# Can AI Conduct Autonomous Scientific Research? Case Studies on Two Real-World Tasks

**DOI:** 10.64898/2026.01.05.697809

**Authors:** Shreyansh Agrawal, Harsh B. Anadkat, Kiran K. Athimoolam, Harsh Bhardwaj, Trishul Chowdhury, Shengtao Gao, Purva K. Kamat, Vishwadeepsinh Makwana, Mohammed H. Shariff, Amitesh Badkul, Lei Xie, Anton V. Sinitskiy

## Abstract

Recent advances in artificial intelligence (AI) have prompted claims about autonomous “AI scientists,” yet systematic evaluations of these capabilities remain scarce. This exploratory study investigates whether current AI frameworks can execute scientific research tasks beyond isolated demonstrations. We tested eight open-source AI frameworks (Agent Laboratory, AutoGen, BabyAGI, GPT Researcher, MOOSE-Chem2, SciAgents, SciMON, and Virtual Lab) on two tasks that aimed to reproduce research on algorithm development from recent papers in uncertainty quantification and protein interaction discovery. In our evaluation, no framework completed a full research cycle from literature understanding through computational execution to validated results and scientific paper writing. While all systems showed competence in conceptual tasks such as planning and summarization, they consistently failed at robust implementation. Every framework produced sophisticated hallucinations. Deployment proved demanding, requiring substantial debugging and technical expertise, which undermines common claims about the democratization of science with AI. Despite these limitations, the frameworks showed promise as research assistants for methodological planning and ideation under careful human supervision. Our findings suggest that the explored AI systems cannot yet autonomously conduct scientific research, but may provide real value for specific subtasks within the research workflow. We offer preliminary observations to help researchers and developers better understand the gap between advertised and actual capabilities of AI in science.

## Introduction

Recent advances in large language models (LLMs) and agentic artificial intelligence (AI) frameworks have fueled ambitious claims in popular press articles, industrial whitepapers, and academic publications about autonomous AI scientists capable of independently designing experiments, running analyses, and writing publishable manuscripts.^1-29^ Despite this rhetoric, there is little systematic independent evaluation of whether existing frameworks can in fact execute a complete and novel scientific project outside of curated demonstrations.^28,30-34^ Most evidence consists of isolated success stories, often reported by the developers of the systems themselves, with limited visibility into failures, boundaries, or the degree of human steering involved. This gap between marketing language and empirical assessment complicates the discussion of what current AI can and cannot do in real scientific practice.

In this work we focus on a more modest question: To what extent can contemporary AI frameworks actually contribute to conducting scientific research, in the sense implied by current narratives about AI “co-authors” and “AI collaborators”? Rather than evaluating isolated capabilities, such as code generation or literature search, we explore whether these systems can orchestrate the set of activities that working scientists would recognize as a research project. More specifically, we start from an existing recent research paper and aim to reproduce or extend its core findings with existing advanced AI frameworks. The goal is not to deliver a definitive verdict on autonomy, but to probe how far current frameworks can go when given realistic tasks that require coordination across multiple skills. Addressing this question involves shifting attention from component-level benchmarks to task-level case studies that more closely approximate the complexity and open-endedness of real scientific work.

To make this question operational, we adopt a working notion of a full research cycle as encompassing a series of essential capabilities: understanding existing scientific literature with sufficient depth to identify knowledge gaps, formulating testable hypotheses and experimental plans, implementing computational methods through correct code, executing experiments with proper controls and parameter settings, obtaining and validating numerical results, interpreting findings within theoretical frameworks and identifying limitations, and producing clear scientific writing that accurately represents methods and results. (We limit the scope of considered scientific research in this work to method development and computational experiments, while wet-lab experiments and scientific research for new discovery of “unknown unknown” were not included due to limited resources available to us on this project.) By this standard, an AI framework that can only propose a high-level plan or generate a draft manuscript without underlying computations does not complete a research cycle, even if its outputs look superficially similar to scientific work. In our analysis, we therefore treat the cycle as a holistic object: we pay attention to partial successes on individual stages, while using the ability to traverse all stages with limited human intervention as a tentative reference point for discussions of “autonomy.”

The landscape of AI frameworks that aspire to support such cycles is heterogeneous and shaped by differing incentives. Many of the most visible systems are commercial and closed-source, exposing only high-level APIs or hosted interfaces, and providing marketing materials that emphasize success while revealing less about internal limitations or failure rates. These systems have incentives to highlight autonomy, and external researchers are typically unable to inspect their internal decision processes or reproduce internal benchmarks. In contrast, open-source frameworks are often developed in academic groups or grant-funded consortia, with code and configuration made publicly available. While these frameworks allow independent inspection to a much greater degree, they frequently suffer from patchy documentation and limited maintenance once initial funding or volunteer labor is exhausted. Our case studies are intended to provide an initial, comparative view of what these different types of systems currently appear to deliver in real scientific contexts, without assuming that they represent the full design space or ultimate performance ceiling.

It is also important to situate any assessment of AI frameworks for science within the broader context of the reproducibility crisis.^35-38^ Across many fields, a substantial fraction of published results have proven difficult or impossible to reproduce, due to incomplete reporting, fragile code and data pipelines, undisclosed selection effects, and sometimes questionable research practices. LLMs add a new layer of risk: they are prone to hallucination, confidently asserting methods, parameter settings, data sources, and numerical results that have no basis in actual computation. When such models are embedded in multi-agent research frameworks, hallucinations can become structurally integrated into experimental plans, analysis scripts, and narrative write-ups, making it difficult for human users to distinguish genuine computation from plausible fabrication. Our analysis is therefore partly exploratory: we seek to understand how these known tendencies of AI systems interact with existing reproducibility challenges, and under what conditions they might mitigate or exacerbate them.

Beyond methodological concerns, there are broader sociotechnical stakes that motivate a careful, exploratory evaluation. Governments, funding agencies, and institutional leaders increasingly view AI as a lever to accelerate scientific and technological innovation. If widely publicized expectations about autonomous AI research systems are not met in practice, the resulting disappointment could fuel public skepticism, reallocation of funding away from both AI and basic science, and a loss of trust among high-level decision makers. Historical episodes of overpromising in AI have contributed to “AI winters,” in which investment and enthusiasm collapsed following unmet expectations. While our case studies cannot predict whether similar dynamics will recur, they are intended to offer a more grounded picture of current capabilities, which may help calibrate expectations and inform more measured discussions among stakeholders.

Independent evaluations introduce important qualifications about the reliability and generality of current AI systems. For example, critical reviews of AI in drug discovery and related fields^9,32,39-44^ highlight pervasive benchmark issues, such as data leakage, distributional mismatch, small and biased datasets, which can inflate reported performance and make apparent breakthroughs difficult to translate into real-world settings. Despite substantial investment and widely publicized timelines from hit identification to early-stage trials, there is still limited evidence that AI-designed candidates improve clinical or industrial success rates compared to traditional approaches.^45^ In parallel, small-scale academic evaluations report that many published “AI scientist” systems do not reproduce their advertised results on new tasks, often require extensive manual debugging, and sometimes hallucinate numerical outcomes or experimental details when underlying computations are absent.^31,34,46^ Together, these independent perspectives suggest that while AI is already useful as a component in scientific pipelines, particularly for ideation, design, and data curation, its current performance and reliability appear to fall short of the fully autonomous, domain-general research agents often implied in promotional narratives.

Existing evaluations of AI in scientific and technical domains, while valuable, thus leave several aspects of the overall picture underexplored. Coding benchmarks typically measure the ability of models to solve small, self-contained problems or to complete functions within well-specified unit tests; they do not assess integration across literature, code, data, and analysis. Other studies showcase impressive case studies in which a system appears to autonomously discover a molecule, optimize a catalyst, or design an experiment, but often provide limited information about failure cases, baseline comparisons, or the extent of human steering. In many instances, it remains unclear whether highlighted successes are representative of typical system behavior or are the result of cherry-picking favorable runs and tasks. Furthermore, benchmark tasks are frequently far simpler and more constrained than contemporary research problems, and they rarely require the kind of interpretive judgment and domain-specific reasoning that real scientific projects demand. Our aim is not to replace these evaluations, but to complement them with a different lens.

Against this backdrop, we present a multi-framework, case-study-based exploration centered on two real-life scientific tasks drawn directly from very recent (and therefore, not included in LLM training sets) papers^47,48^ and implemented using real-world code and data. Across these tasks, we examine the behavior of eight distinct AI frameworks: Agent Laboratory,^24^ AutoGen,^49^ BabyAGI,^50^ GPT Researcher,^51^ MOOSE-Chem2,^52^ SciAgents,^4^ SciMON,^53^ and Virtual Lab.^26^ Each framework is tasked with attempting to reproduce or extend the core results of a target paper, starting from publicly available materials and subject to realistic constraints on compute and human time. We document not only successes and failures at each stage of the research cycle, but also the human interventions required, the kinds of hallucinations observed, the stability of the tools under repeated use, and the extent to which outputs might be misleading if taken at face value. This design is intended to provide an initial comparative picture of how current frameworks behave on complex, end-to-end scientific tasks, recognizing that any conclusions are necessarily conditional on the specific domains, papers, and configurations we study.

On the basis of these case studies, the paper offers tentative, evidence-informed observations and recommendations for both users and developers of AI research frameworks, with the goal of helping to navigate the gap between current promise and practice. For researchers in the natural sciences, we highlight specific situations in which existing systems already appear to provide substantial value, for example, accelerating literature review, structuring experimental plans, and generating candidate analyses, while also pointing to stages of the research cycle where our observations suggest that careful human oversight remains crucial.

For developers and policymakers, we discuss possible design principles and reporting practices that could make future claims about “autonomous” research more interpretable, including more systematic documentation of failure modes, explicit treatment of hallucination risks, and better support for traceable, reproducible computation. Rather than offering a definitive verdict on “AI scientists,” our aim is to contribute an early, empirically grounded perspective on how current frameworks interact with real scientific workflows, and to outline questions that future work will need to address.

## Results

### None of the evaluated AI frameworks completed a full scientific research cycle

Despite claims of autonomous scientific research, our empirical evaluation revealed that all explored frameworks fundamentally operated as conceptual planning and text generation tools rather than scientific executors (Fig. 1). The frameworks demonstrated strong performance in understanding papers and generating research plans but uniformly failed at the critical transition from planning to implementation. No framework successfully executed code to produce genuine computational results, with only Agent Laboratory sometimes attempting code execution. The disconnect between advertised capabilities and actual performance was most pronounced in frameworks claiming full research reproduction. For example, Agent Laboratory claimed being “capable of completing the entire research process”^24^ but generated complete manuscripts with entirely fabricated results, and Virtual Lab, claimed that it “can rapidly make an impactful, real-world scientific discovery”, only produced code templates and method outlines but could not execute them and required advanced manual debugging.^26^ Only two frameworks (SciMON and MOOSE-Chem2) performed as advertised, but notably these were the only frameworks that explicitly limited their scope to hypothesis generation and conceptual ideation rather than claiming execution capabilities.

**Fig 1.**
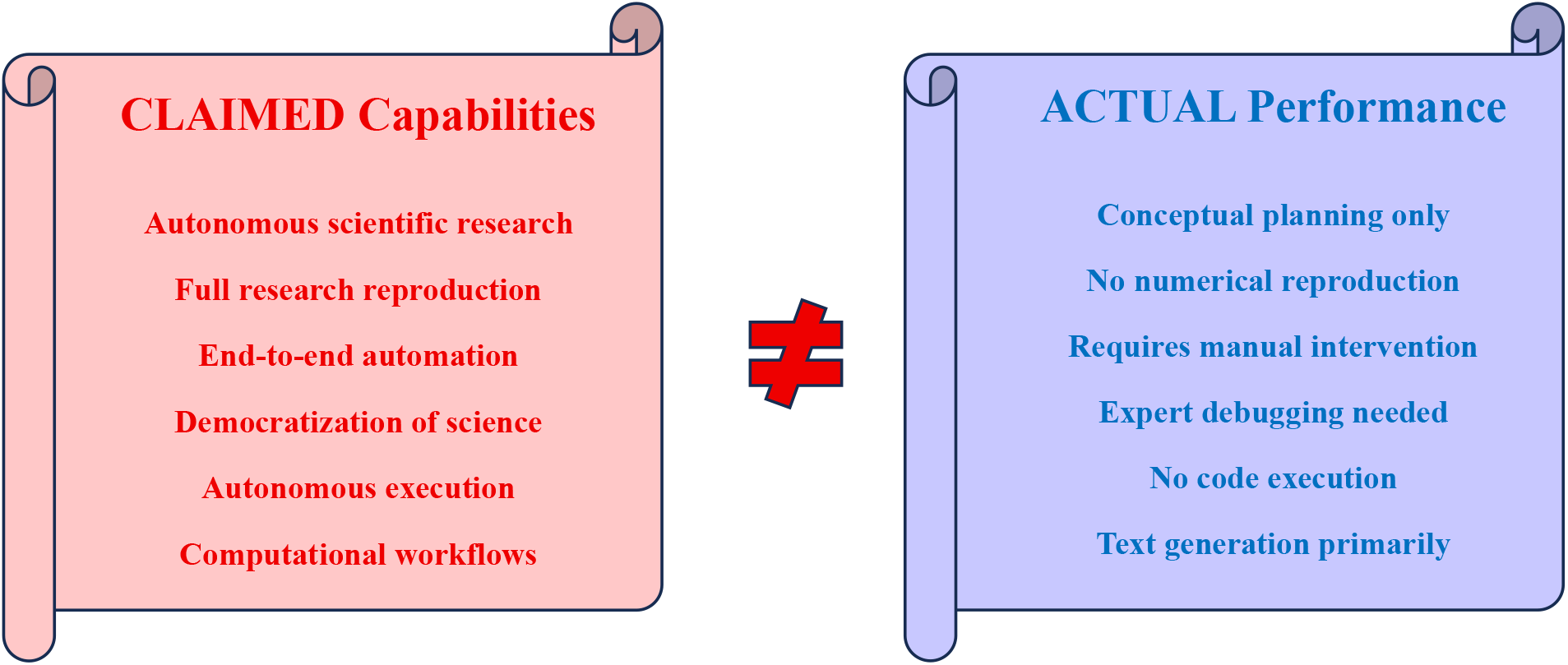
Gap Between Claims and Reality

### All frameworks severely hallucinated scientific methods, data, and results

The hallucination patterns observed were sophisticated enough to potentially deceive peer reviewers, with fabricated outputs including specific numerical values within plausible ranges, error margins suggesting statistical rigor, and technical terminology deployed correctly within context (Table 1). For example, Agent Laboratory produced complete fabrications including “>99% predictive accuracy” or FRET efficiency measurements, and MOOSE-Chem2 suggested maintaining constant temperature, ionic strength, and other experimental conditions for a purely computational project. The multi-agent frameworks exhibited particularly concerning patterns of circular hallucination, where agents would assume other agents had computed results that never existed, creating cascading fabrications that appeared validated through inter-agent consensus. AutoGen agents generated “collective confidence scores” and “inter-agent validation correlations” that seemed to represent rigorous cross-validation but were entirely fictional. Even frameworks that attempted to constrain hallucination through prompt engineering failed to eliminate it entirely: BabyAGI continued to imply structural validation and specific performance improvements even when explicitly instructed to avoid numerical claims, while SciMON generated “novelty scores” and “conceptual coherence metrics” that could easily be misinterpreted as empirical findings. The sophistication of these hallucinations, including dataset-specific metrics, thermodynamically plausible values, and convergence points at reasonable iteration counts, demonstrates that current AI systems can generate scientifically plausible but entirely fictional results that would require advanced domain expertise to identify as fabrications.

**Table 1.**
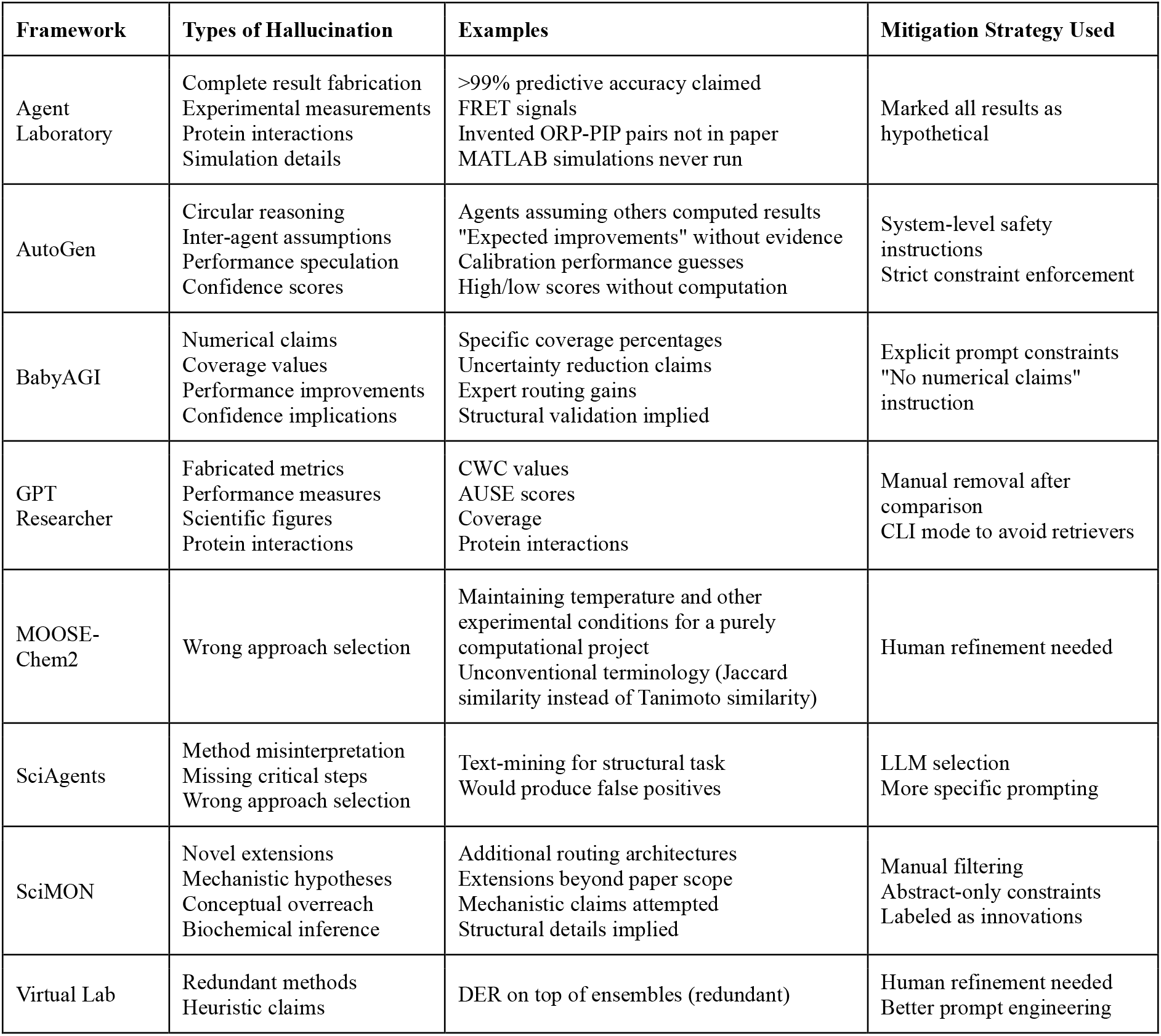
Hallucination Patterns Observed.

### Every AI framework required substantial manual debugging during deployment

None of the frameworks functioned as documented, with setup times ranging from 35 minutes for MOOSE-Chem2 to 6-8 hours for GPT Researcher, and every system encountering critical errors not mentioned in their documentation. Common issues included missing API keys that caused 401 authorization failures (GPT Researcher), undocumented dependencies (BabyAGI required sqlalchemy and flask, AutoGen needed specific PYTHONPATH configuration), and fundamental architectural problems (assertion errors for num_hierarchy parameters in MOOSE-Chem2, Python execution failures when encountering shell syntax in Agent Laboratory). The deployment challenges were particularly severe with the use of HPC resources. SciAgents on HPC required 7-8 hours of debugging including manual Ollama installation, CUDA module conflicts, and GPU detection failures that required extracting entire archive directories and carefully managing library paths. Even after resolving installation issues, runtime problems persisted: AutoGen fell into conversation loop stalls requiring manual stopping conditions, BabyAGI failed when executed outside the repository root directory, and the dependency of Virtual Lab on external tools created cascading configuration requirements. The level of technical expertise required for deployment (including understanding of Python environments, API configurations, HPC job scheduling, CUDA drivers, and framework-specific architectures) directly contradicts claims^13,19,26,54^ about democratization of science and accessibility to non-experts. These deployment challenges suggest that even if the frameworks could perform as advertised, their practical utility would be severely limited by the advanced engineering expertise required for basic operation.

### Case study 1: Uncertainty Quantification Task

The recent paper by Badkul and Xie^47^ proposed an expert-routed conformal prediction method for uncertainty quantification in protein-ligand binding affinity prediction under out-of-distribution (OOD) shift, requiring implementation of mixture-of-experts architecture, split-conformal calibration, and evaluation metrics including Coverage-Width Criterion (CWC), Prediction Interval Coverage Probability (PICP), and Area Under the Size-Error curve (AUSE). Across all frameworks, none successfully reproduced the quantitative results or implemented the core algorithm (Table 2). GPT Researcher produced thorough conceptual descriptions of uncertainty quantification methods and outlined the method implementation but failed to calculate any of the reported metrics, instead generating illustrative values that appeared scientific but lacked computational backing. BabyAGI captured the methodological architecture including the train-calibrate-test pipeline and expert routing logic but could not access the ChEMBL31 dataset or execute the conformal calibration procedures. The more sophisticated frameworks attempted complex but flawed approaches. Virtual Lab proposed combining Deep Evidential Regression with deep ensembles and heteroscedastic heads, effectively stacking several epistemic-uncertainty mechanisms on top of each other. This design suggests a partial misunderstanding of uncertainty decomposition: deep ensembles already provide an epistemic signal, while heteroscedastic heads target aleatoric noise, so adding DER on top introduces redundancy and extra compute without a clear role for each component. In addition, distribution shift was handled via heuristic OOD-based interval inflation rather than a clearly specified shift-aware conformal method. The results from SciAgents varied dramatically by underlying LLM, with Qwen3 proposing Monte Carlo Dropout and Gemini suggesting deep ensembles, both different from the original paper expert-routing approach. Even when frameworks correctly identified key concepts like conformal prediction and expert routing, they could not translate these into executable code or produce the empirical coverage guarantees that define successful uncertainty quantification.

**Table 2.** Framework Performance on Case Studies.

| Framework | Case Study 1: Uncertainty Quantification | Case Study 2: ORP-PIP Interaction Discovery |
| --- | --- | --- |
| GPT Researcher | ⊕ Conceptual description of uncertainty methods<br>⊖ No CWC, coverage, or AUSE calculations<br>⊖ Hallucinated performance metrics<br>⊖ No code execution, numerical results fabricated | ⊕ Described computational procedures<br>⊖ No AlphaFold access or PDB data<br>⊖ No structural predictions or modeling<br>⊖ Invented protein interactions without computation |
| BabyAGI | ⊕ Captured methodological architecture and logic<br>⊕ Pipeline structure with expert routing concepts<br>⊖ No empirical results or dataset access<br>⊖ Fabricated coverage values when unconstrained | ⊕ Conceptual workflow paralleling study<br>⊕ Abstract association tables generated<br>⊖ No AlphaFold computations possible<br>⊖ No structural validation or predictions |
| AutoGen | ⊕ Multi-perspective methodological analysis<br>⊕ Identified calibration assumptions<br>⊖ No quantitative reproduction<br>⊖ Agents assumed others computed non-existent data | ⊕ Collaborative workflow reconstruction<br>⊕ Conceptual prioritization logic<br>⊖ No computational biology capabilities<br>⊖ Hallucinated confidence scores between agents |
| SciMON | ⊕ Well-structured idea cards for each component<br>⊕ Novel architectural extensions proposed<br>⊖ No implementation or code generation<br>⊖ Cannot produce numerical results | ⊕ Structured hypotheses and themes<br>⊕ Conceptual interaction categorization<br>⊖ No structural predictions attempted<br>⊖ Extensions misinterpretable as actual findings |
| MOOSE-Chem2 | ⊕ Detailed MPNN architecture blueprint<br>⊕ CQR model plan with pinball loss<br>⊖ No actual model training or code execution | N/A |
| Agent Laboratory | ⊕ Generated complete manuscript format<br>⊖ Attempted code execution (Python runtime errors)<br>⊖ No numerical results produced<br>⊖ Hallucinated measurements reported in generated manuscript | ⊕ Generated complete manuscript format<br>⊖ Fabricated >99% accuracy claims<br>⊖ Invented FRET signals and biosensor data<br>⊖ Code execution attempts failed with Python errors |
| SciAgents (Qwen3) | ⊕ GNN with Monte Carlo Dropout plan<br>⊖ No implementation or execution<br>⊖ No actual uncertainty intervals computed | ⊖ Completely wrong approach (text-mining)<br>⊖ Would find only known interactions<br>⊖ Missed structural biology context entirely |
| SciAgents (Gemini) | ⊕ Deep Ensemble of GNNs proposed<br>⊖ No implementation provided<br>⊖ No actual results generated | ⊕ Correct AlphaFold-Multimer approach<br>⊖ Missed critical co-localization filter<br>⊖ Would produce many false positives |
| Virtual Lab | ⊕ Deep Evidential Regression + Ensembles proposed<br>⊖ Redundant epistemic signals (DER on ensembles)<br>⊖ Under-specified shift-aware calibration<br>⊖ Post-hoc OOD width inflation instead of WCP | ⊕ Planned AlphaFold-based screening<br>⊖ Could not execute all-versus-all runs<br>⊖ Missing binaries/dependencies<br>⊖ No structural predictions produced |

### Case study 2: Protein Interaction Discovery

The recent study by Dall’Armellina et al.^48^ used AlphaFold structural predictions with new generation confidence scores to identify interactions between OSBP-related proteins (ORPs) and phosphoinositide 3-, 4-, and 5-phosphatases (PIPs) at membrane contact sites, requiring both computational structure prediction and biological validation through co-localization filtering. The AI attempts to reproduce this work revealed fundamental limitations in handling domain-specific scientific workflows (Table 2). GPT Researcher described computational procedures and potential ORP-PIP interaction couples but lacked access to AlphaFold binaries, PDB databases, or GPU-accelerated structure prediction tools, resulting in confident but unsubstantiated protein interaction claims. Agent Laboratory generated a complete manuscript including fabricated Förster resonance energy transfer (FRET) signals, biosensor outputs, and specific interaction pairs that were not present in the original paper, while also inventing MATLAB simulation details for analyses never performed. SciAgents with Qwen3 completely misunderstood the task, proposing a text-mining pipeline to extract known interactions from literature rather than discovering new ones through structural prediction, while the same framework with Gemini correctly identified AlphaFold-Multimer as the appropriate tool but critically missed the co-localization filtering step that prevents biologically impossible false positives. This omission would have resulted in a candidate list dominated by spurious interactions between proteins that never encounter each other in cellular contexts. AutoGen produced conceptual ORP-PIP association tables through multi-agent collaboration, but the agents hallucinated confidence scores and implied structural validation without any actual AlphaFold computations. The AI failures in this case study were particularly revealing of their limitations in biological domains: inability to access specialized scientific software, lack of understanding of biological constraints and cellular organization, and tendency to generate plausible-sounding protein names and interactions without grounding in structural or experimental data. Even Virtual Lab, which advertised integration with AlphaFold and other structural biology tools, could not execute the basic all-versus-all screening of approximately 240 ORP-PIP pairs that formed the foundation of the original study.

### AI frameworks demonstrated value as research assistants for conceptual reasoning and methodological planning

On the positive side, all AI frameworks excelled at decomposing complex scientific problems into structured workflows, with BabyAGI producing particularly clear hierarchical task breakdowns including tables, checklists, and evaluation schemas that accurately captured the logical flow of both case studies. SciMON generated the most academically coherent outputs through its idea card format, producing publication-ready conceptual frameworks that included inspiration sources, novelty assessments, evaluation plans, and explicit failure mode analyses that could meaningfully guide human researchers in experimental design. AutoGen provided uniquely valuable multi-perspective methodological interpretations, with different AI agents identifying complementary assumptions and potential improvements, for instance, correctly recognizing that conformal prediction requires independent and identically distributed conditions and that scaffold-based splits violate these assumptions in ways requiring specialized handling. Hierarchical hypothesis refinement in MOOSE-Chem2 demonstrated sophisticated understanding of experimental design, correctly identifying the need for fingerprints, UniRep and ESM protein embeddings, and pinball loss functions for quantile regression, even though it could not implement these components. The frameworks also showed remarkable ability to synthesize methodological insights across domains. Virtual Lab proposed to combine ensemble methods with conformal prediction, which correctly identified the complementary nature of parametric and non-parametric uncertainty quantification approaches. Perhaps most significantly, when frameworks operated within their true capabilities, as evidenced by SciMON and MOOSE-Chem2 performing exactly as advertised when limited to hypothesis generation, they produced valuable scientific insights including novel architectural proposals, identification of methodological assumptions that might otherwise be overlooked, and structured research plans that could accelerate the initial phases of scientific investigation. These successes suggest that AI frameworks, when properly scoped and with transparent limitations, can serve as powerful tools for literature synthesis, hypothesis generation, and methodological planning, functioning as research assistants that augment rather than replace human scientific expertise.

## Discussion

Our study is not intended as another benchmark but as a multi-framework, case-based stress test targeting modern, non-trivial scientific tasks that resemble the work scientists actually do. Rather than asking whether models can pass coding quizzes or synthetic exam-style benchmarks, we asked several AI frameworks to grapple with recent, complex research papers that were unlikely to be represented in their training data, together with real codebases, datasets, and deployment constraints. This design exposed not only performance ceilings but also the detailed failure modes that arise when ambitious claims about autonomous research are confronted with end-to-end workflows: brittle tool integrations, mismatched assumptions about data availability, and subtle conceptual errors that would be invisible on toy tasks. In this sense, our results complement existing leaderboard-style evaluations by shifting attention from scalar accuracy metrics to qualitative patterns across planning, execution, error recovery, and interaction with human users.

Viewed through this lens, our findings suggest that current frameworks are best understood as automating specific research sub-tasks, especially planning, drafting, and, to a lesser extent, coding, rather than executing full research cycles. Across both case studies, systems were most reliable when asked to decompose problems into subtasks, outline methodological pipelines, draft literature reviews, or propose code skeletons. They repeatedly faltered at the transitions that matter for genuine scientific contribution: from pseudocode to executable code, from hypothetical metrics to computed results, from textual description of structural biology workflows to actual runs of software like AlphaFold, or from uncertainty quantification plans to calibrated prediction intervals with demonstrable coverage properties. Even where code generation occurred, it was rarely integrated into reproducible pipelines with correct data handling, evaluation, and logging. Taken together, these observations suggest that, at present, autonomous scientific research in the strong sense is better interpreted as “semi-automated support for selected phases of scientific work,” and that claims of full-cycle automation require substantial qualification.

The contrast between closed-source and open-source frameworks in our study also has implications for how the field should be evaluated. Several of the most ambitious claims came from commercial, closed systems that expose only high-level interfaces and marketing materials, making it difficult to verify internal behavior, reproduce advertised results, or diagnose failures. In contrast, the open-source frameworks, though often poorly documented and fragile, allow one to inspect configuration files, trace tool calls, and identify where design decisions or implementation bugs led to specific failure modes. This transparency enables more fine-grained, critical evaluation and suggests that systematic study of open frameworks may provide a useful baseline for assessing claims about closed systems with similar functionality, even if the latter remain partially opaque.

Hallucination in this setting is not merely a nuisance but an epistemic and institutional risk, because fabricated methods, data, and results can easily be laundered into scientific discourse via AI-generated papers. Our case studies show that modern AI systems can generate highly realistic experimental narratives, including plausible parameter settings, confidence intervals, and domain-appropriate jargon, with no underlying computation. In multi-agent settings, these hallucinations can be reinforced through apparent “consensus” between agents, making them harder to detect. If such outputs are incorporated into manuscripts or presentations without rigorous checking, they risk polluting the literature with non-existent results, misleading reviewers and readers, and distorting downstream meta-analyses or policy decisions. This dynamic is particularly concerning in high-stakes fields like biomedicine, where fabricated performance numbers, false-positive interaction predictions, or misrepresented uncertainty estimates could influence resource allocation, trial design, or safety assessments.

These risks are intertwined with a structural tension between optimizing for impressive demos and building robust, repeatable deployments in real labs. Many AI frameworks are showcased through carefully curated examples where data are pre-cleaned, tools are pre-configured, and runs are cherry-picked for success. Our attempts to deploy the same systems on independent tasks revealed that small deviations from the demo environment, such as different cluster setups, missing binaries, or slight changes in task framing, often led to cascading failures. In several cases, framework design choices appeared to prioritize visually compelling artifacts (such as automatically generated “papers” with figures and tables) over traceability and reproducibility of the underlying computations. This tension is not unique to our case studies; it reflects broader incentives in the AI ecosystem, where demo-ready prototypes can attract attention and funding even if they do not yet support the reliability and observability needed for everyday scientific use.

The difficulty of deployment we observed even among expert users calls into question narratives about the imminent “democratization of science” via autonomous AI. Running these frameworks required substantial expertise in software engineering, HPC environments, dependency management, and domain-specific tooling, as well as patience for debugging vague errors and undocumented behaviors. If such friction is typical, then the benefits of these tools may accrue primarily to well-resourced groups with strong engineering support, rather than to small laboratories or individual researchers. Paradoxically, this suggests that the current generation of “autonomous research” systems may *increase* dependence on specialist infrastructure and skills, rather than lowering the barriers to participation in scientific inquiry.

Against this backdrop, hype and overselling around “AI scientists” pose a real threat to the credibility of the broader AI and scientific communities. Overstated claims about autonomy, robustness, and impact can generate short-term excitement but risk eroding trust when they collide with the realities of fragile deployments, hallucinated results, and modest practical gains. Our case studies do not imply that AI is useless for science; on the contrary, they reveal clear value in literature synthesis, hypothesis generation, and experimental planning. But they do suggest that the gap between headline narratives and everyday performance remains substantial. If this gap is not acknowledged and managed, there is a danger that stakeholders, including funders, institutional leaders, and the public, will become increasingly skeptical of AI-based approaches, potentially undermining support for genuinely promising work and reinforcing cycles of boom-and-bust investment.

These observations motivate several tentative recommendations for users, funders, and developers. For scientific users, our results suggest treating current AI frameworks as powerful but fallible assistants: tools for brainstorming, structuring projects, and surfacing edge cases, rather than authoritative sources of results. Rigorous validation, including independent recomputation of any numerical claims and explicit documentation of which steps were human-versus AI-executed, should be standard practice. For funders and institutional decision makers, supporting independent evaluations, open benchmarks that reflect realistic workflows, and infrastructure for reproducible experiments may be more impactful than sponsoring isolated flagship demos. Developers, both commercial and academic, could mitigate risks by clearly stating the intended scope and limitations of their systems, surfacing uncertainty about their own outputs, and providing mechanisms for tracing how particular results were produced (for example, through execution logs and environment captures). Across all stakeholders, aligning incentives toward reliability, transparency, and reproducibility, rather than headline-grabbing claims of autonomy, appears crucial.

Our findings also have pedagogical and training implications. In classroom settings, we observed that agentic frameworks can serve as catalysts for critical discussion, helping students articulate research questions, enumerate design choices, and anticipate failure modes. Used carefully, these tools can support instruction in scientific reasoning by providing draft plans and arguments that students must then scrutinize, correct, and ground in data. At the same time, their propensity for producing confident but incorrect content underscores the need to teach “AI literacy” as part of scientific education: students should be trained to treat AI outputs as hypotheses to be tested rather than authorities to be trusted. Assignments that explicitly require tracing the provenance of results, identifying hallucinated claims, and contrasting AI-generated plans with manual baselines may help cultivate the skepticism and methodological rigor needed to work productively with these systems.

Looking ahead, we see several directions for future work. Methodologically, larger and more diverse case studies across knowledge domains would help clarify how general our observations are, and whether some scientific areas are more amenable to partial automation than others. On the systems side, there is scope for developing AI frameworks that more tightly integrate language models with containerized, reproducible execution environments, robust tool-checking, and explicit separation between speculative and computed statements. Evaluation-wise, new benchmarks that capture not only task success but also deployment friction, hallucination risk, and human time saved would offer a more nuanced picture of system utility. Finally, interdisciplinary collaborations between AI researchers, domain scientists, and science studies scholars could yield richer accounts of how these tools reshape scientific practice, including their impact on credit, labor, and institutional structures.

In light of our results, how should we answer the initial question, “Can AI conduct autonomous scientific research?” Our case studies suggest that, at present, the answer depends crucially on how “conduct” and “autonomous” are understood. If the phrase is taken to mean end-to-end execution of full research cycles, with reliable code, real experiments, validated results, and accountable documentation, then our observations indicate that current frameworks do not yet achieve this standard, and claims to the contrary should be treated with caution. If, however, we interpret the question more narrowly, as asking whether AI can meaningfully contribute to scientific inquiry by automating parts of the workflow, then our findings are more optimistic: today’s systems can already accelerate literature review, scaffold experimental design, and enrich methodological reflection when used under careful human oversight. In this sense, the more productive reformulation may not be whether AI can replace scientists, but how AI can be designed and governed to act as a robust, transparent, and accountable collaborator within scientific research.

## Methods

### Selection of AI frameworks

We selected eight AI frameworks for evaluation, based on three criteria: public code availability as open-source repositories on GitHub with sufficient documentation to attempt deployment, explicit claims about scientific research capabilities in their repositories or associated papers, and representation of different architectural approaches to research automation. The AI frameworks spanned diverse technical approaches and development contexts despite their common open-source nature: BabyAGI^50^ and GPT Researcher^51^ represented task-decomposition systems with claims of autonomous research capabilities; AutoGen,^49^ Agent Laboratory,^24^ and Virtual Lab^26^ exemplified multi-agent architectures where specialized agents collaborate on different aspects of research; SciAgents,^4^ SciMON^53^ and MOOSE-Chem2^52^ focused on hypothesis generation and scientific ideation (in the case of MOOSE-Chem2, in a narrow domain field). While all frameworks provided source code access, they varied significantly in their origins and maintenance patterns: some emerged from well-funded research groups or companies that open-sourced their tools, others from academic grant-funded projects, and several from individual developers or small collaborations. This variation in development context, despite universal code availability, influenced documentation quality, maintenance frequency, and the gap between advertised capabilities and actual functionality. We did not include frameworks lacking scientific focus, those requiring additional proprietary components not freely available, and repositories where documentation was insufficient to attempt deployment despite code availability. The final selection represented frameworks developed over the last few years, with repository activity ranging from actively maintained projects with recent commits to apparently abandoned codebases that had not been updated for months.

### Target tasks and source papers

We selected two contemporary scientific papers published after the knowledge cutoff dates of most language models to prevent memorization-based performance and ensure tasks represented genuine research challenges rather than retrieval exercises. The first task involved reproducing the main results from an arXiv manuscript “Adaptive Individual Uncertainty under Out-Of-Distribution Shift with Expert-Routed Conformal Prediction” by Amitesh Badkul and Lei Xie, published on Oct 17, 2025,^47^ which proposed a sophisticated uncertainty quantification approach for protein-ligand binding affinity prediction using mixture-of-experts architecture, conformal prediction, and specialized handling of scaffold-based dataset splits to address distribution shift in drug discovery applications. This task required understanding of advanced statistical methods including conformal prediction theory, implementation of complex neural architectures with expert routing mechanisms, handling of chemical datasets with proper scaffold splitting, and computation of specialized metrics including Coverage-Width Criterion, Prediction Interval Coverage Probability, and Area Under the Size-Error curve. The second task was based on the bioRxiv manuscript “AlphaFold-driven discovery of ORP-PIP phosphatase interactions using new generation confidence scores” by Dall’Armellina, Urbé, and Rigden, published on Sept 15, 2025^48^, which identified novel protein-protein interactions between ORP and PIP proteins through structural prediction and biological validation. This task demanded different capabilities: running AlphaFold-Multimer for structure prediction, understanding protein families and cellular localization, implementing biological filters to remove false positives, and interpreting structural confidence metrics. Both papers are very recent and technically non-trivial, making it unlikely that complete solutions were present in model training data and ensuring that the tasks stress real research workflows rather than memorization. Together, these tasks tested both methodological sophistication in machine learning and domain expertise in structural biology, requiring integration of conceptual understanding with practical implementation.

### Experimental design and operator role

Evaluations were performed by human operators (master’s level students in an advanced AI course at Northeastern University), who acted as expert but non-adversarial users of each framework. Each operator was responsible for installing one or more AI frameworks from public repositories, configuring them according to the official documentation, and running them on the target tasks under realistic resource constraints. Each framework was provided with a brief natural language description of the reproduction task. Operators interacted with the systems through their recommended interfaces (command-line tools or Jupyter notebooks), starting from a shared template prompt tailored to each paper and then iteratively refining prompts only as needed to clarify the task or to enforce constraints such as “do not invent numerical results.” They were allowed to fix environment and configuration issues (for example, missing Python dependencies, incorrect API keys, HPC job submission errors, and path problems). Importantly, operators were allowed to fix environment and configuration issues (e.g., dependencies, API keys) but did not provide substantive domain-level corrections to algorithms or tune models to achieve claimed numerical results. Operators performed first-pass screening to flag and exclude obviously fabricated outputs before synthesis, but they did not supply domain-expert re-analyses or re-computations of numerical claims or verify the correctness of generated results during execution. Each framework was run in its recommended environment, with some deployed locally on laptops (BabyAGI, AutoGen, SciMON, MOOSE-Chem2, Agent Laboratory, Virtual Lab), others on HPC clusters (GPT Researcher, SciAgents). During each run, operators kept informal logs of prompts, configuration changes, errors encountered, time spent on setup versus interaction, and the AI framework intermediate and final outputs, and they performed a first-pass screening to flag obviously hallucinated claims before passing materials to the synthesis stage.

### Analysis approach and limitations

Our analysis employed multiple complementary methods to assess framework performance across the complete research cycle. We evaluated task completion by examining whether frameworks successfully traversed all stages from literature understanding through result generation, documenting partial successes at individual stages while treating the research cycle holistically. Hallucination detection involved comparing all numerical claims, methodological descriptions, and scientific assertions against ground truth from the original papers and known scientific principles, categorizing hallucinations by type and sophistication level. Deployment burden was quantified through setup time, debugging requirements, undocumented dependencies, and the level of technical expertise required for basic operation. We analyzed the gap between advertised and actual capabilities by systematically comparing repository README files, documentation, and associated papers with observed performance. The evaluation has several important limitations that constrain generalizability: we tested only two papers from specific domains (structural biology and machine learning), leaving uncertain how frameworks might perform on wet-lab experiments or other computational fields; the rapid development pace means frameworks may have improved since evaluation; our sample size of selected AI frameworks, while substantial given the deployment effort required, may not represent the full spectrum of available approaches; the expertise level and debugging persistence of the human operator likely influenced successful deployment rates; and our focus on reproduction tasks may not capture the potential of these AI frameworks for novel discovery or creative research. As a result, our findings should be interpreted as an initial, context-dependent map of behaviors under realistic but constrained conditions, rather than as definitive rankings or generalizable performance guarantees. Despite these limitations, our evaluation provides the first extensive, comparative assessment of open-source AI frameworks on complete, contemporary scientific tasks, offering empirical grounding for discussions about autonomous AI research that have previously relied primarily on promotional materials or isolated examples.

## Supplementary Information

**Table S1.**
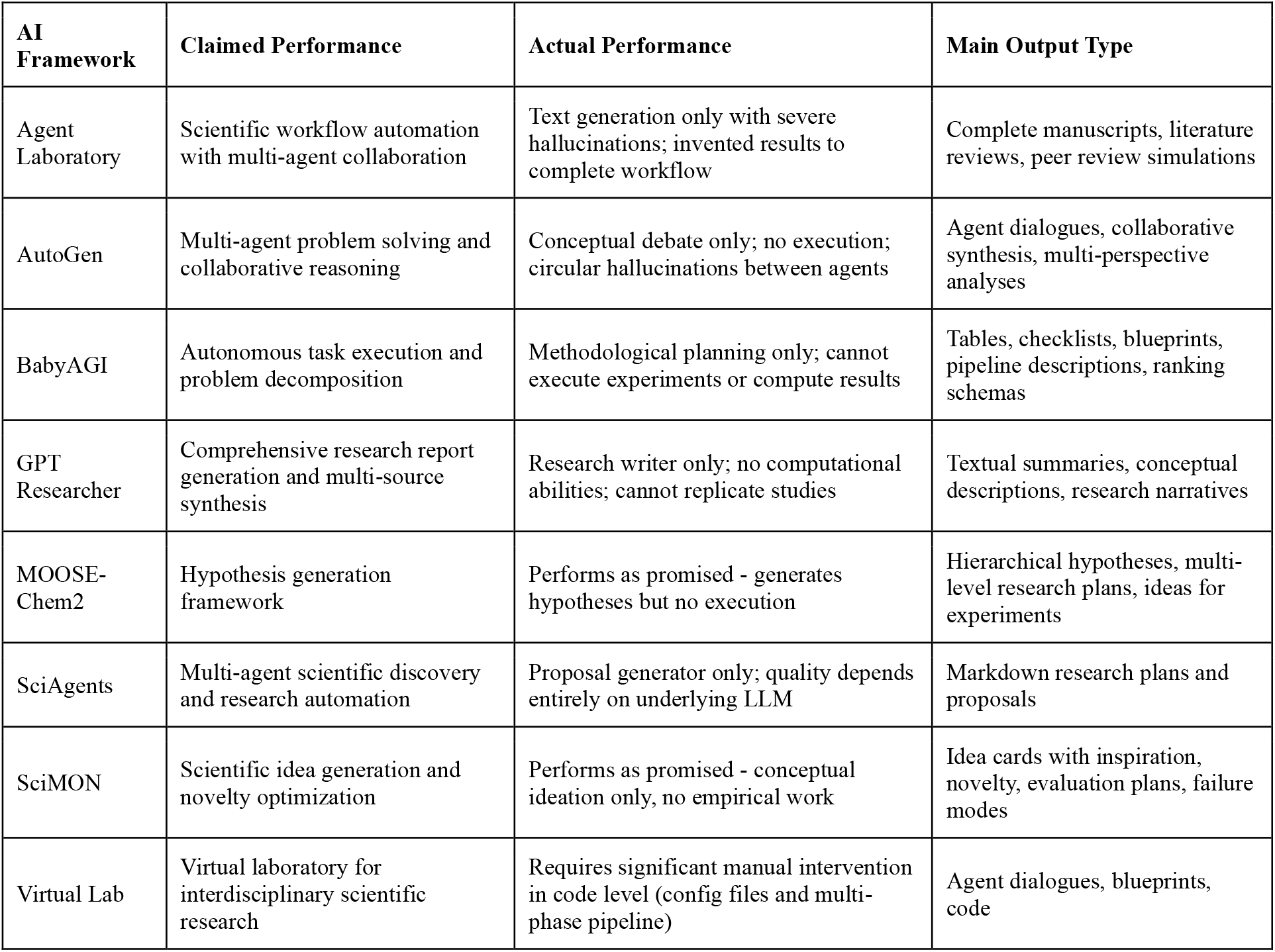
Summary of AI Frameworks Evaluated.

| AI Framework | Claimed Performance | Actual Performance | Main Output Type |
| --- | --- | --- | --- |
| Agent Laboratory | Scientific workflow automation with multi-agent collaboration | Text generation only with severe hallucinations; invented results to complete workflow | Complete manuscripts, literature reviews, peer review simulations |
| AutoGen | Multi-agent problem solving and collaborative reasoning | Conceptual debate only; no execution; circular hallucinations between agents | Agent dialogues, collaborative synthesis, multi-perspective analyses |
| BabyAGI | Autonomous task execution and problem decomposition | Methodological planning only; cannot execute experiments or compute results | Tables, checklists, blueprints, pipeline descriptions, ranking schemas |
| GPT Researcher | Comprehensive research report generation and multi-source synthesis | Research writer only; no computational abilities; cannot replicate studies | Textual summaries, conceptual descriptions, research narratives |
| MOOSE-Chem2 | Hypothesis generation framework | Performs as promised - generates hypotheses but no execution | Hierarchical hypotheses, multi-level research plans, ideas for experiments |
| SciAgents | Multi-agent scientific discovery and research automation | Proposal generator only; quality depends entirely on underlying LLM | Markdown research plans and proposals |
| SciMON | Scientific idea generation and novelty optimization | Performs as promised - conceptual ideation only, no empirical work | Idea cards with inspiration, novelty, evaluation plans, failure modes |
| Virtual Lab | Virtual laboratory for interdisciplinary scientific research | Requires significant manual intervention in code level (config files and multi-phase pipeline) | Agent dialogues, blueprints, code |

**Table S2.**
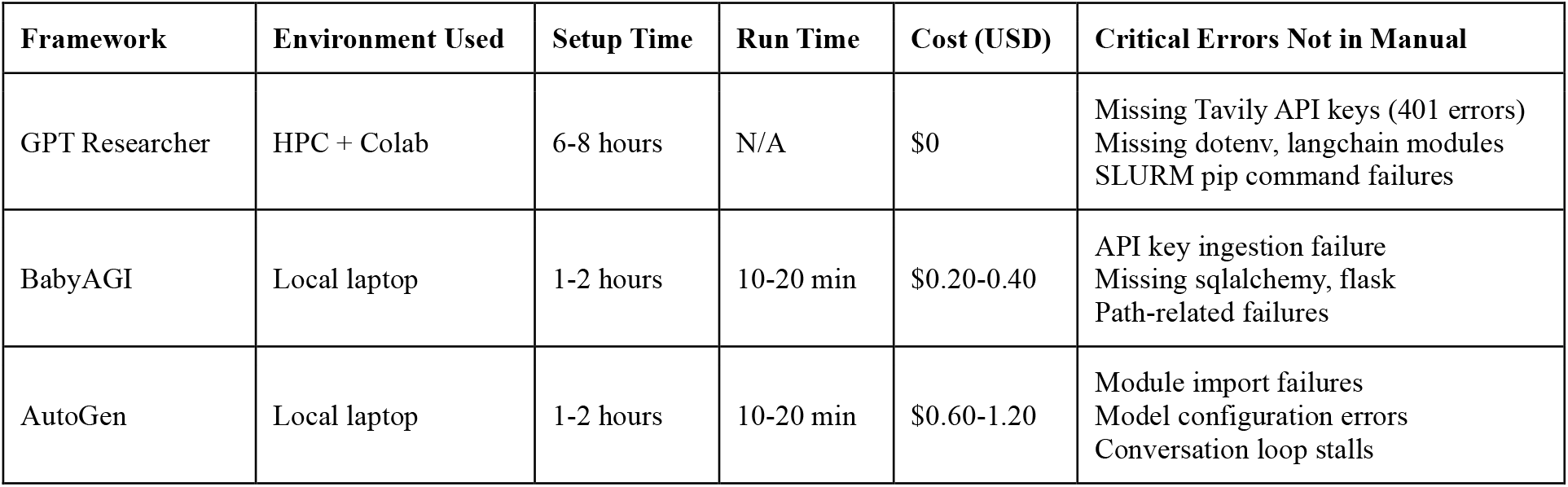
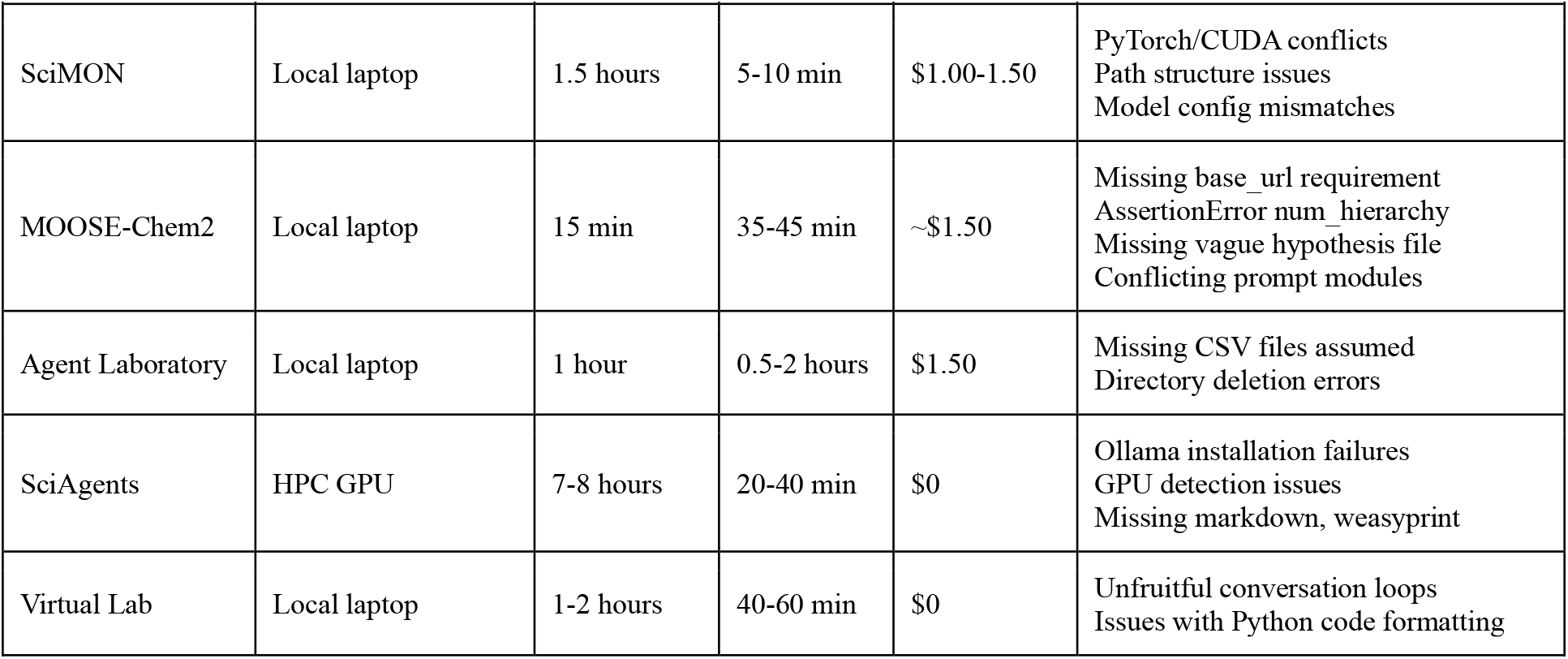
Deployment Metrics and Technical Challenges.

**Table S3.** Representative Hallucination Examples.

| Framework | Hallucinated Output | Why It Would Fool Reviewers |
| --- | --- | --- |
| GPT Researcher | Fabricated uncertainty metrics (CWC, AUSE, coverage)<br>Confident protein-interaction claims without any computation. | Numbers looking realistic<br>Domain-specific terminology<br>Sounded empirically calculated |
| MOOSE-Chem2 | Maintaining temperature and other experimental conditions for a purely computational project | Technical terminology correct<br>Looks like a solid research plan<br>Appears empirically determined |
| Agent Laboratory | More than 99% predictive accuracy<br>FRET signals<br>Invented ORP-PIP interaction pairs<br>Fake MATLAB simulation details | Experimental validation implied |
| AutoGen | ORP-PIP interaction table | Reasonable improvement margin<br>Just above target threshold<br>High but believable correlation<br>Acceptable drift range |
| Virtual Lab | Partial misunderstanding of uncertainty decomposition | Appears empirically determined<br>Reasonable variance estimate<br>Better than baseline expected<br>Physical parameter range |
| BabyAGI | Narrative summary claiming multi-agent agreement | Multi-agent agreement suggests validation<br>Different perspectives converging<br>Technical terminology correct<br>Specific architecture choice |
| SciAgents | Text-mining pipeline for structural discovery | Correct technical terms<br>Looks like a solid research plan |

